# The role of MYB21 in Arabidopsis: transcending flower-specific functions to vegetative tissue

**DOI:** 10.64898/2026.03.10.710835

**Authors:** Khansa Mekkaoui, Lisa Thuy Linh Nguyen, Olivia Putri Herdani, Selma Gago Zachert, Stefan Mielke, Ranjit Baral, Gerd Hause, Ivan F. Acosta, Debora Gasperini, Bettina Hause

**Affiliations:** Department of Cell and Metabolic Biology, Leibniz Institute of Plant Biochemistry, Weinberg 3, D-06120 Halle/Saale, Germany; Martin Luther University Halle-Wittenberg, Institute of Biochemistry and Biotechnology, Charles Tanford Protein Center, Kurt-Mothes-Str. 3a, D-06120 Halle/Saale, Germany; Department of Molecular Signal Processing, Leibniz Institute of Plant Biochemistry, Weinberg 3, D-06120 Halle/Saale, Germany; Martin Luther University Halle-Wittenberg, Biozentrum, Core Facility Microscopy, Weinbergweg 22, D-06120 Halle/Saale, Germany; Max Planck Institute for Plant Breeding Research, Carl-von-Linné-Weg 10, D-50829 Köln, Germany

**Keywords:** *Arabidopsis thaliana*, *Botrytis cinerea*, COI1-dependent signaling, defense responses, insect herbivory, jasmonic acid (JA), R2R3-MYB transcription factor, vegetative development

## Abstract

Jasmonic acid (JA) and its derivatives are lipid-derived phytohormones that coordinate plant growth, development, and stress responses through the bioactive conjugate jasmonoyl-isoleucine (JA-Ile). Their role in reproductive development is well established, particularly in stamen maturation through the R2R3-MYB transcription factor MYB21, which has been considered largely flower-specific. Here, we reveal a previously unrecognized role of MYB21 in vegetative tissues of *Arabidopsis thaliana*. Although basal *MYB21* transcript levels in leaves are extremely low and spatially restricted, wounding and exogenous hormone applications induced *MYB21* transcription in a JA- and COI1-dependent manner. Transcriptional GUS reporter analyses showed localized *MYB21* promoter activity in specialized epidermal cells, including trichomes, hydathodes, and in the vasculature at wound sites. Functional characterization using the *myb21-5* mutant indicated roles in germination and vegetative growth, partially phenocopying JA-insensitive mutants despite unaltered JA biosynthesis and signaling. Transcriptome profiling further revealed changes in expression of genes involved in lignin biosynthesis, light-harvesting complex components, cytokinin pathways, and defense-related responses, consistent with reduced resistance of *myb21-5* to insect herbivory and infection by *Botrytis cinerea*. Together, these findings identify MYB21 as a JA-responsive regulator of growth and defense in seedlings and leaves, extending its function beyond reproductive development.

**Highlight:** The transcription factor MYB21, previously considered flower-specific, is induced by jasmonate signaling in Arabidopsis leaves and seedlings, where it regulates growth and defense responses, thereby expanding its role beyond reproductive development.

## Introduction

Jasmonates (JAs) are lipid-derived phytohormones with well-established functions in several plant processes, including root growth (Staswick *et al*., 1992), reproductive development (Park *et al*., 2002; Stintzi and Browse, 2000), abiotic stress responses (Wang *et al*., 2021b) as well as in plant wounding response (Delessert *et al*., 2004; Howe, 2004) and defense against herbivorous insects and pathogen attacks (Howe and Jander, 2008; Li *et al*., 2022). To coordinate plant growth and stress responses, JAs and particularly the bioactive jasmonoyl-isoleucine (JA-Ile), primarily act as signaling molecules mediating transcriptional reprogramming (Fonseca *et al*., 2009). The process of JA-Ile signaling in plants involves a series of molecular events initiated by its perception by the SCF-COI1-JAZ co-receptor complex (Sheard *et al*., 2010), which leads to the degradation of the JASMONATE ZIM-DOMAIN (JAZ) repressors (Chini *et al*., 2007; Thines *et al*., 2007; Yan *et al*., 2007) and subsequent activation of downstream JA-Ile responses (Pauwels *et al*., 2008).

A common feature among mutants deficient in jasmonic acid (JA) biosynthesis and perception in *Arabidopsis thaliana* is male sterility, characterized by defects in stamen filament elongation, anther dehiscence and pollen maturation (Browse, 2009). Except for the JA-insensitive mutant, *coronatine insensitive 1* (*coi1)*, exogenous JA applications rescue the male fertility of these mutants (Sanders *et al*., 2000; Stintzi and Browse, 2000). The role of JA in stamen development was elucidated through transcriptional profiling of JA-treated stamens from the JA biosynthesis mutant deficient in OXO-PHYTODIENOIC ACID-REDUCTASE 3 (*opr3*), and led to the identification of the JA-responsive MYB21 and MYB24 transcription factors as regulators of this process (Mandaokar *et al*., 2006). Both transcription factors belong to the clade 19 of the R2R3-MYB family (Dubos *et al*., 2010), and are interactors of several JAZ repressors (Song *et al*., 2011). While the *myb24* mutant has no obvious defects, the *myb21* mutant shows a male sterility phenotype, visible by shortened anther filaments and delayed anther dehiscence (Mandaokar *et al*., 2006). In response to jasmonate, MYB21 is proposed to mediate stamen and pollen maturation in coordination with MYB24 and MYB108 at stage twelve of flower development (Mandaokar *et al*., 2006; Reeves *et al*., 2012; Song *et al*., 2011). However, starting from stage thirteen, it shifts its role to suppress JA biosynthesis, thereby initiating a negative feedback loop that arrests flower maturation processes (Reeves *et al*., 2012). MYB21 also regulates flavanols biosynthesis in the flower stamens together with the transcription factors MYB24 and MYB57 (Zhang *et al*., 2021).

*MYB21* exhibits tissue-specific expression predominantly in flower organs, such as sepals, petals, stamen filaments, and carpels (Reeves *et al*., 2012). Its orthologs in tomato and other plant species function also in flowers (Schubert *et al*., 2019; Wang *et al*., 2021a; Yang *et al*., 2020), confirming its flower-specific role. The detection of *MYB21* transcripts in seedlings of the *constitutive photomorphogenic 1* (*cop1)* mutant indicated that *MYB21* expression in Arabidopsis vegetative tissues is normally repressed by the light signaling component COP1. Consistently, ectopic expression of *MYB21* induced developmental anomalies resembling those observed in the *cop1* mutant, including dwarfism, small leaves, and seedling lethality (Shin *et al*., 2002; Song *et al*., 2011). These observations suggest that *MYB21* expression is repressed in vegetative tissues and any possible function of MYB21 in vegetative processes has been so far neglected.

Here, we report that *MYB21* is transcriptionally induced in vegetative tissues of *A. thaliana* (whole seedlings and rosettes of adult plants) in response to wounding, in a JA- and COI1-dependent manner, despite its very low to undetectable expression levels under basal conditions. Using a *MYB21_pro_:GUSPlus* reporter line and in agreement with its transcript levels, we show that *MYB21* expression in the leaf is restricted to few epidermal cells at basal conditions, which extend to specialized epidermal cells such as trichomes after leaf wounding and exogenous hormone applications. Mechanical injury also triggers *MYB21* expression in the leaf vasculature, but only near the wound site. Analysis of the *myb21-5* mutant (Reeves *et al*., 2012) reveals defects in vegetative growth along with decreased resistance to insect feeding and *Botrytis* infection. These phenotypes closely resemble those observed in JA-insensitive or deficient mutants, even though *myb21-5* does not show detectable alterations in JA biosynthesis and signaling. Together these findings point to a JA-mediated function of MYB21 in vegetative growth and defense responses, despite a spatially restricted expression in specific vegetative cell types.

## Material and methods

### Plant material and growth conditions

*Arabidopsis thaliana* wild type (ecotype Col-0), *myb21-5* (Reeves *et al*., 2012), *coi1-16* (Ellis and Turner, 2002), and *aos/dde2-2* (von Malek *et al*., 2002) seeds as well seeds of transgenic lines expressing *MYB21_pro_:GUSPlus* (in Col-0) or *MYB21_pro_:MYB21* (in *myb21-5,* (Schubert *et al*., 2019) were surface sterilized in 4% bleach and stratified for three days in the dark at 4°C. Seedlings were grown either in liquid Murashige and Skoog (MS) medium (pH 5.7, 1% sucrose) or on solid MS medium (1% agar), while adult plants were individually cultivated in pots containing steam-sterilized clay, coir fiber, and vermiculite. Growth was conducted in Phytocabinets (Percival Scientific, www.percivalscientific.com) under short-day conditions, with a 10/14-hour light/dark cycle, temperatures of 21/19°C, 65% relative humidity and a light intensity of 120 µE m^-2^ s^-1^.

### Generation of transgenic line expressing *MYB21_pro_:GUSPlus*

The 7281-bp regulatory region upstream of the AtMYB21 coding sequence has been previously reported (Schubert *et al*., 2019). We cloned it via Gateway recombination in front of the reporter gene *GUSPlus*^TM^ (Broothaerts *et al*., 2005) in a binary destination vector containing the seed selection marker *OLE1*pro-*tagRFP* (Shimada *et al*., 2010). The resulting pMP72 plasmid was stably transformed in Arabidopsis via the flower dipping method (Clough and Bent, 1998). Twenty-one stable lines were propagated and shown to display similar expression patterns in maturing stamens. A single line was further advanced to T3 and used for all experiments presented here.

### Wounding and hormone treatment

For gene expression analyses and hormone measurements, wounding was performed in the morning (9-11 am) either by squeezing grouped seedlings under sterile conditions or pinching the middle of the mature leaf using haemostats with serrated teeth. One hour after wounding, grouped seedlings were harvested, tissue-dried and snap-frozen in liquid nitrogen, while mature leaves or full rosettes were cut off and snap-frozen. For local wound experiments, the upper half of the leaf was pinched with a haemostat, and one-hour post-wounding, both leaf halves – the upper half (wounded area) and lower half (adjacent area) – were separately harvested. MeJA treatment was carried out in the morning by adding a 50 µM solution to either the solid or liquid MS media of the seedlings. Adult plants were sprayed with 100 µM MeJA three times to ensure full coverage of the rosettes.

### RNA isolation, RT-qPCR and transcriptome analyses

Total RNA was isolated from frozen material using the RNeasy Plant Mini Kit (Qiagen, 74904) and genomic DNA was removed using the DNA-free™ DNA Removal Kit (Invitrogen, AM1906). Reverse transcription was performed using the RevertAid H Minus First Strand cDNA Synthesis Kit (Thermo Scientific™, K1631) with Oligo(dT)_18_ primers (Thermo Scientific™, SO131). The generated cDNA was used as template in 10 µL reaction mix containing 1x EvaGreen QPCR Mix II (Bio&Sell, BS76.580.0200) and 0.2 µM of forward and reverse primers (Supplemental Table 1). qPCR was run on a CFX Connect RT-PCR Detection System (Bio-Rad Laboratories) with the following protocol: denaturation (95°C for 15 min), amplification (40 cycles of 95°C for 15 s and 60°C for 30 s) and melt curve analysis (95°C for 10 s, 65°C heating up to 95°C with a heating rate of 0.05°C s^−1^). Primers were designed using the Primer3plus software (www.primer3plus.com) and checked on the Beacon Designer™ software (www.premierbiosoft.com). Gene expression was calculated in relation to the *PROTEIN PHOSPHATASE 2A SUBUNIT A3 (PP2AA3)* (Czechowski *et al*., 2005) using the 2^-ΔCT^ method (Schmittgen and Livak, 2008) and included biological triplicates.

For transcriptome analyses, RNA isolated from non-wounded seedlings of wild type and *myb21-5* was subjected to RNA quality and integrity assessment on an Agilent 2100 Bioanalyzer system (Agilent Technologies, www.agilent.com) using the RNA 6000 Nano Kit for standard RNA sensitivity (Agilent, #5067-1511). Three biological replicates of each condition were submitted to Novogene (www.novogene.com) for mRNA paired-end short-read sequencing (150 bp length) on an Illumina NovaSeq 6000 Sequencing System and bioinformatics analysis according to their pipeline. Mapping of reads, determination of gene expression levels, and generation of gene expression heatmaps, gene clustering, and Gene Ontology (GO) enrichment analysis were done as described (Mekkaoui *et al*., 2025).

### Phytohormone measurements

Measurements of OPDA, JA, and JA-Ile were performed using a standardized Ultra-performance liquid chromatography–tandem Mass Spectrometry (UPLC– MS/MS)-based method (Balcke *et al*., 2012). 50 mg of frozen powdered tissue were extracted with 500 µl 100% LC-MS methanol supplemented with 50 ng of [^2^H_5_]OPDA, [^2^H_6_]JA, and [^2^H_2_]JA-Ile each as internal standards. After solid-phase extraction on HR-XC (Chromabond, Macherey-Nagel, www.mn-net.com), 10 µl of the eluate were analyzed via UPLC–MS/MS, and analyte content was determined relative to the internal standard peak heights.

### Histochemical detection of GUS activity

Four-week-old rosettes from the *MYB21_pro_:GUSPlus* reporter line were either wounded by a haemostat or sprayed with 100 µM MeJA. One-hour post-treatment, leaves were detached from the rosettes, fixed in 90% acetone for one hour and subjected to GUS staining according to (Stenzel *et al*., 2012). Chlorophyll removal was achieved by incubation series in ethanol. Open flower buds at stage 13 were GUS-stained as described by (Mudunkothge and Krizek, 2014). Imaging was performed in chloral hydrate:glycerol:water (8:2:1, v/v/v).

### Root length and leaf area measurements

Sterile seeds of Col-0 and *myb21-5* were placed 1 cm apart on solid MS medium and grown vertically for seven days for root measurements, and horizontally for two weeks for leaf measurements. Images were captured using a Stemi 2000-C stereo microscope (Carl Zeiss) equipped with an AxioCam MRc5 camera (Carl Zeiss) and processed by ImageJ software for measurements. Primary root length was measured in ten experiments, each with five seedlings, while leaf area and trichome number were measured in three experiments, each with ten seedlings.

### Seeds germination assay

Col-0 and *myb21-5* seeds from the same year’s stocks were sown on solid MS medium and imaged at 24, 30, 48, 54 and 72 hours at the onset of daylight. The designation of the different stages of seed germination and development relied on the classification as published by (Maia *et al*., 2011) for the initial stages and (Silva *et al*., 2017) for the later stages.

### Determination of cell size in leaves

Small pieces taken from the same position of leaf 13 from fully developed rosettes from Col-0 and *myb21-5* were fixed in 3% (v/v) sodium cacodylate-buffered glutardialdehyde (pH 7.2), dehydrated in an ethanol series, and embedded in epoxy resin (Spurr, 1969). Semi-thin sections (1 µm thickness) were stained with toluidine blue. Micrographs were taken using a Zeiss ‘AxioImager’ microscope (Zeiss) equipped with an AxioCam (Zeiss). The size of all parenchyma cells of a cross section was determined using the measurement tool from Photoshop® 2020 (Adobe Systems). Measurements were performed from four different biological replicates each.

### Herbivore and pathogen assays

Herbivory bioassays were performed as described in (Mielke and Gasperini, 2020). In brief, plants were grown for 33 days in short-day photoperiod (8 h light / 16 h dark) at 21 °C, 60 % relative humidity and a light intensity of 100 µE/m2/s. 11 plants per genotype were transferred into 20 x 30 x 20 cm Plexiglas boxes and three newly hatched *Spodoptera littoralis* larvae were placed on each plant. Boxes were brought back to the growth chamber and larvae were allowed to feed for a maximum of 10 days. Surviving larvae were collected and individually weighed.

To multiply spores from *B. cinerea* strain B05.10 canned peach halves were inoculated with and incubated in a sealed transparent container for 5 days in a plant chamber (CLF Plant Climatics, Wertingen, Germany) under a 12 h light/ 12 h dark photoperiod at 22° C/20° C. Spores were collected from the surface of the inoculated peach and placed into a Falcon tube containing 20 mL B5-Glc medium (3.16 g/L solid Gamborg B5 Medium with vitamins (Duchefa) and 2% glucose (w/v)) followed by filtering through a double layer of Miracloth (Merck KGaG, Darmstadt, Germany). After adjusting spore suspension to 5×10^5^ spores/mL, simultaneous spore germination was initiated by adding phosphate buffer at pH 6.4 to a final concentration of 10 mM and incubation at RT for 1 h. Leaves No. 8, 9, 10 and 11 from 4-week-old *A. thaliana* plants of Col-0 wild type, *myb21-5* mutant, and MYB21-complemented *myb21*-mutant were drop-inoculated with 10 µL of spore-suspension per leaf. Plants were kept in trays to avoid evaporation and to maintain 100% humidity. Pictures from the leaves were taken at 48 h post-inoculation and the lesion area was determined with ImageJ (Schneider *et al*., 2012).

### Statistical analysis

The statistical analyses applied to the different datasets as indicated in the figures were performed using GraphPad Prism (www.graphpad.com).

## Results

### *MYB21* expression is induced by wounding and MeJA in vegetative tissues

MYB21 has been considered a flower-specific transcription factor in *A. thaliana* and other plant species, with expression and characterized functions confined to floral organs and plant fertility, as reported by previous studies (Mandaokar *et al*., 2006; Reeves *et al*., 2012; Shin *et al*., 2002; Song *et al*., 2011). Consistent with these studies, *MYB21* transcript levels are notably high in Arabidopsis flower organs, but are either undetectable or remain at very low levels in vegetative tissues such as young seedlings and mature leaves (CoNeKT database; https://conekt.sbs.ntu.edu.sg/) (Fig. 1A). Recent transcriptomic datasets of wounded ten-day-old seedlings and leaves of adult plants from Col-0, showed a strong induction of *MYB21* transcripts, with log_2_ fold change values above 5 and 8, respectively (Supplemental Table S2), despite generally low expression levels. This observation suggested possible participation of MYB21 in the wound response and possible function in vegetative processes besides its crucial function in plant fertility. First, the kinetics of *MYB21* transcript accumulation upon wounding was examined by RT-qPCR in Col-0 seedlings and showed a maximum of transcript accumulation at 1-3 hour after wounding which declined over the subsequent 24 hours (Fig. 1B). This expression pattern parallelled that of the JA marker gene *JASMONATE-ZIM-DOMAIN PROTEIN 10* (*JAZ10*), suggesting that the wound-induced accumulation of *MYB21* transcripts might be related to JA signaling (Fig. 1B). Putative JA-regulation of *MYB21’s* expression in vegetative tissues was investigated in rosette leaves treated with 100 µM JA methyl ester (MeJA) which is de-esterified in planta into JA (Fig. 1C), and showed that *MYB21,* similarly to *JAZ10,* was responsive to MeJA in the leaves as its transcripts accumulated significantly at 60 and 90 minutes after treatment (Fig. 1C). To determine the JA-dependence of *MYB21* expression in Col-0 leaves, its accumulation upon wounding and MeJA-treatment was determined in the JA-deficient mutant *allene oxide synthase* (*aos*; Von Malek et al. 2002) and the JA-insensitive mutant *coronatine insensitive 1* (*coi1-16;* Ellis and Turner 2002) (Fig. 1D-E). *MYB21* transcript accumulation did not occur in the absence of JA-Ile synthesis or perception, as shown for the *aos* and *coi1-16* mutants, respectively, compared to the wild type (Fig. 1D). Consistently, *MYB21* transcript accumulation in response to MeJA was detected in wild-type plants and the *aos* mutant but not in the *coi1-16* mutant (Fig. 1E), indicating that *MYB21* expression in Arabidopsis leaf is both JA- and COI1-dependent. The transcription factor MYB24 has been characterized to function in conjunction with MYB21 during flower maturation, and its gene is also predominantly expressed in flowers (Cheng et al. 2009; Reeves et al. 2012) while almost undetected in leaves. Therefore, its expression in response to wounding and MeJA was analyzed in Col-0 (Supplemental Fig. S1). *MYB24* transcripts accumulated in wounded seedlings of Col-0 and in MeJA-treated mature leaves (Supplemental Fig. S1A-B), and these transcriptional responses occurred in a JA- and COI1-dependent manner (Supplemental Fig. S1C). This suggested that, like MYB21, MYB24 function(s) may not be restricted to fertility and floral organs.

**Fig. 1:**
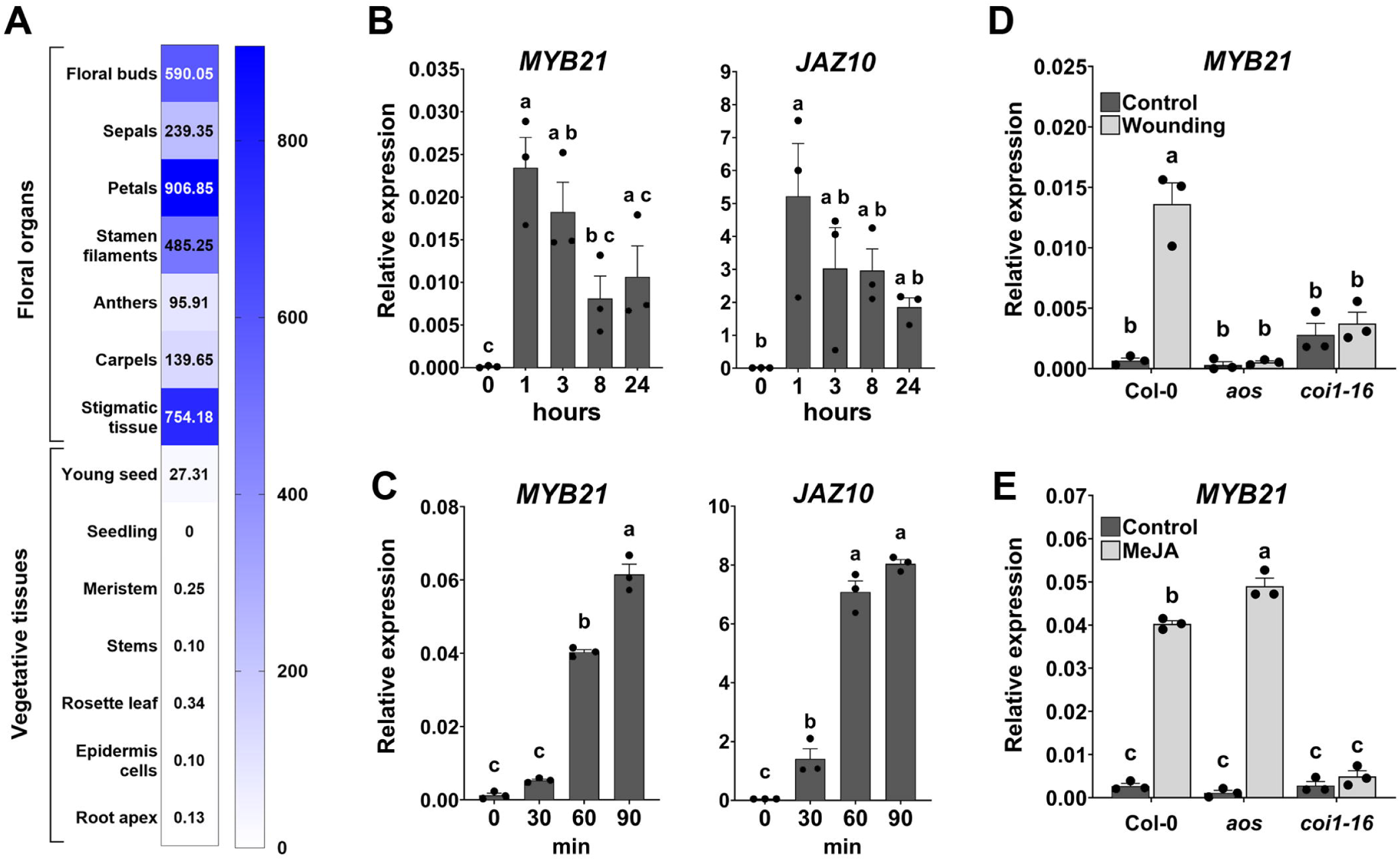
MYB21 transcript accumulation in leaves is wound-induced and JA-dependent. (A) Organ- and tissue-specific expression of *MYB21* in *Arabidopsis thaliana.* The heatmap depicts the expression profiles of *MYB21* in transcripts per million (TPM), sourced from the CoNekT database (https://conekt.sbs.ntu.edu.sg/). Note the main expression of *MYB21* in the floral organs, whereas it is almost absent in the vegetative tissues. (B) Transcript accumulation of *MYB21* and *JAZ10* in 14-day-old seedlings at 0, 1, 3, 8 and 24 h after wounding. (C) Transcript accumulation of *MYB21* and *JAZ10* in rosette leaves at 0, 30, 60, and 90 minutes following spraying with 100 µM MeJA. (D) Transcript level of *MYB21* in rosette leaves from four-week-old plants before (Control) and one hour after wounding (Wounding) in wild type (Col-0), *aos* and *coi1-16*. (E) Transcript level of *MYB21* in rosette leaves from four-week-old plants before (Control) and one hour after spraying with 100 µM MeJA (MeJA) in wild type (Col-0), *aos* and *coi1-16*. Transcript levels were determined by RT-qPCR and normalized to those of *AtPP2A3*. Bars represent means of three biological replicates (single dots; ±SEM). Statistically significant differences among time points (B-C) were calculated using One-Way ANOVA, and among time points and genotypes (D-E) using two-way ANOVA, followed by Tukey-HSD and are indicated by different letters. **Associated supplemental data: Fig. S1**

### *MYB21* promoter shows trichome- and vasculature-specific activity in the leaf

To uncover the expression pattern and tissue-specificity of *MYB21* in the leaf, its promoter activity was determined by GUS assay using a *MYB21*(7kb)*pro::GUSPlus* reporter line. Initial verification of the 7 kb *MYB21* promoter activity in open flower buds at stage 13 of development showed *MYB21* expression in the apical part of the stamens and gynoecium, as well as in the petals and sepals (Supplemental Fig. S2A), which is consistent with previous reports (Reeves et al. 2012). In the absence of a stimulus (referred to as control condition), GUS staining was specific to the leaf hydathode areas and to trichomes and was barely detectable in the leaf blade (Fig. 2A-C). The hydathode- and trichome-specific expression of *MYB21* aligns with its very low transcripts levels in a whole leaf and seedling when determined by RT-qPCR or RNA-seq (Fig. 1). Upon spraying the leaf with 100 µM MeJA, *MYB21* promoter activity was induced in the leaf margin around the entire leaf and in the trichomes (Fig. 2D). *MYB21* promoter showed activity in the basal cells of the trichomes and the epidermal cells surrounding them in response to MeJA, which was not detected at control conditions (Fig. 2D-F). A similar induction of *MYB21* promoter activity in the trichomes and their basal cells occurred at 1 hour after wounding of the leaf with forceps (Fig. 2G-I) and was accompanied by an activity in the leaf midrib vasculature exclusively within the wounded area (Fig. 2H). Cross-sections through the lamina of wounded and control leaves further revealed *MYB21* promoter activity in the vascular bundles and epidermal cells near the wounding site, which were absent under control conditions (Fig. 2J-K). To determine *MYB21* promoter activity in younger plants, wounding was performed on cotyledons and primary leaves of 15-day-old seedlings (Fig. S2). Similar to adult plants, *MYB21* promoter activity was induced by wounding in the vasculature, the leaf margin and the trichomes, while at control condition no expression near the hydathodes was visible indicating a specific expression in mature hydathode areas at later stages (Supplemental Fig. S2B-E).

**Fig. 2:**
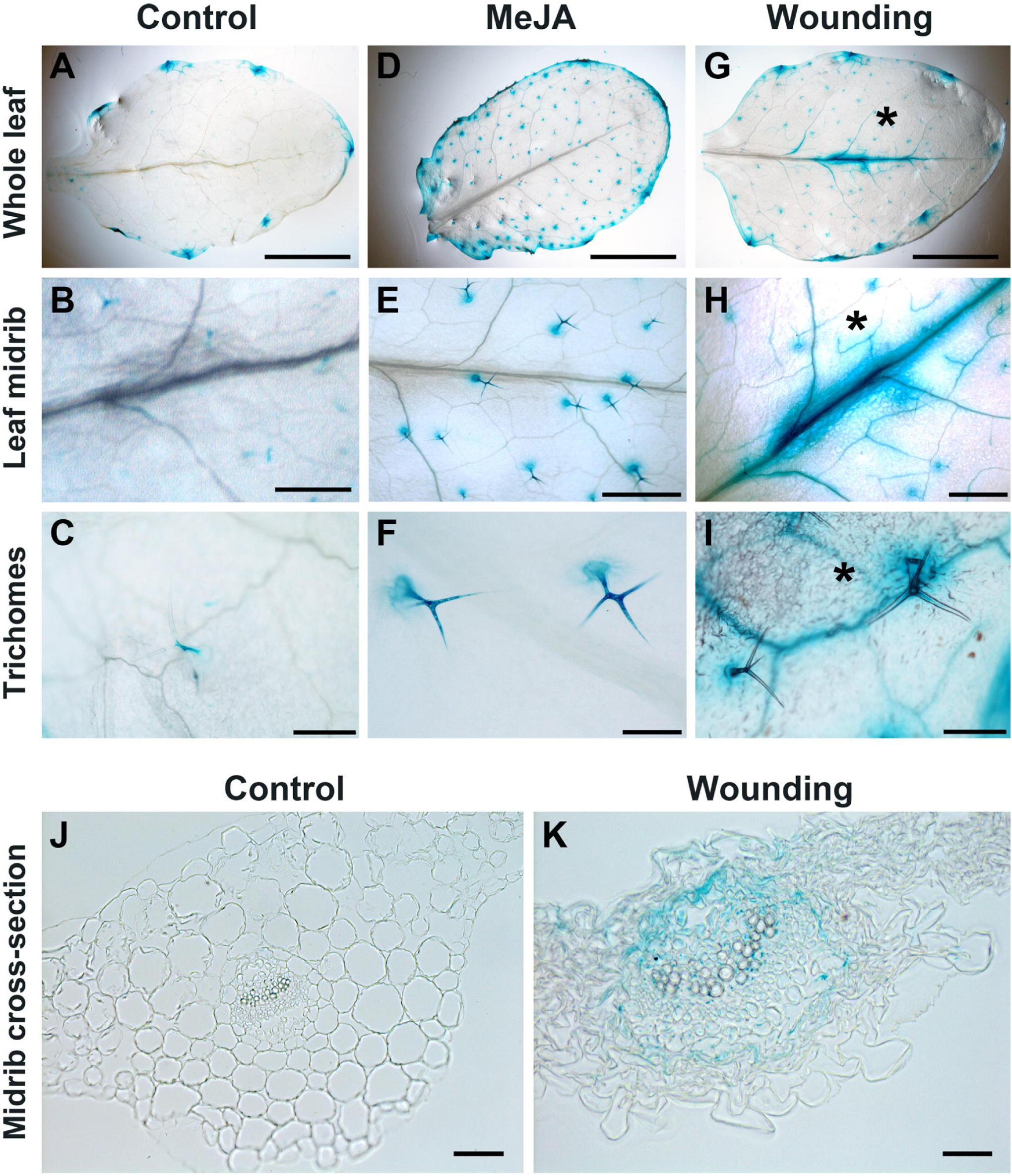
Activity of *MYB21* promoter in Arabidopsis leaves in response to wounding and MeJA treatment. Histochemical detection of GUS in four-week-old leaves from the *MYB21_pro_:GUSPlus* line at control condition (A-C), 1 hour following leaf spraying with 100 µM MeJA (D-F), and 1 hour following wounding of the leaf with haemostat (G-I), including cross-sections of the leaf midrib (J-K). (A, D, G) Whole leaf. (B, E, H) Leaf midrib and vasculature. (C, F, I) Leaf trichomes. (J-K) Cross-sections of the leaf midrib before (J) and at 1 hour after wounding (K). Asterisks indicate the wounded area. Scale bars represent 2 mm (A, D, G), 500 µm (B, E, H), 200 µm (C, F, I) and 50 μm (J, K). **Associated supplemental data: Fig. S2**

### A MYB21-mediated feedback regulation of JA biosynthesis or signaling is not detected in mature leaves and seedlings

Given that in stamen development, MYB21 is proposed to negatively regulate JA biosynthesis in a feedback loop from flower stage 13 onwards (Reeves *et al*., 2012), we investigated whether MYB21 also regulates JA biosynthesis or signaling during the wound response in seedlings using the knock-out mutant *myb21-5* (Reeves *et al*., 2012). In wounded seedlings, *myb21-5* did not statistically differ from the wild-type in transcript levels of JA-responsive genes (Fig. S3A-B), or JA/JA-Ile production (Fig. S3C). Similarly, protein accumulation of the JA-biosynthesis enzymes ALLENE OXIDE CYCLASES (AOCs) which is indicative of JA-positive feedback loop during development (Stenzel *et al*., 2003), did not show differences between *myb21-5* and wild-type (Fig. S3D-E), suggesting that MYB21 does not mediate a feedback regulation of JA biosynthesis or signaling during wounding and at control conditions.

Considering the local and midvein-specific expression of *MYB21* in leaf tissue upon wounding, we next assessed JA-related responses specifically within the wounded area, since localized effects could be masked in whole-leaf analyses (Fig. 3). Additionally, the unwounded area adjacent to the wound site was analyzed, to examine a possible role of MYB21 in the wound response of distant tissues. In line with the GUS assay results, *MYB21* transcript induction occurred predominantly within the wounded area (Fig. S4). Comparing *myb21-5* and wild-type leaves, almost no differences were detected in the local and systemic JA-mediated wound responses, as determined by JA/JA-Ile levels (Fig. 3A) and transcript accumulation of early and late JA marker genes (Fig. 3B-D). An exception to this pattern was the JA-responsive *TERPENE SYNTHASE 4 (TPS4),* which showed elevated transcript levels in the wounded area of the *myb21-5* mutant compared to the wild type (Fig. 3D).

**Fig. 3:**
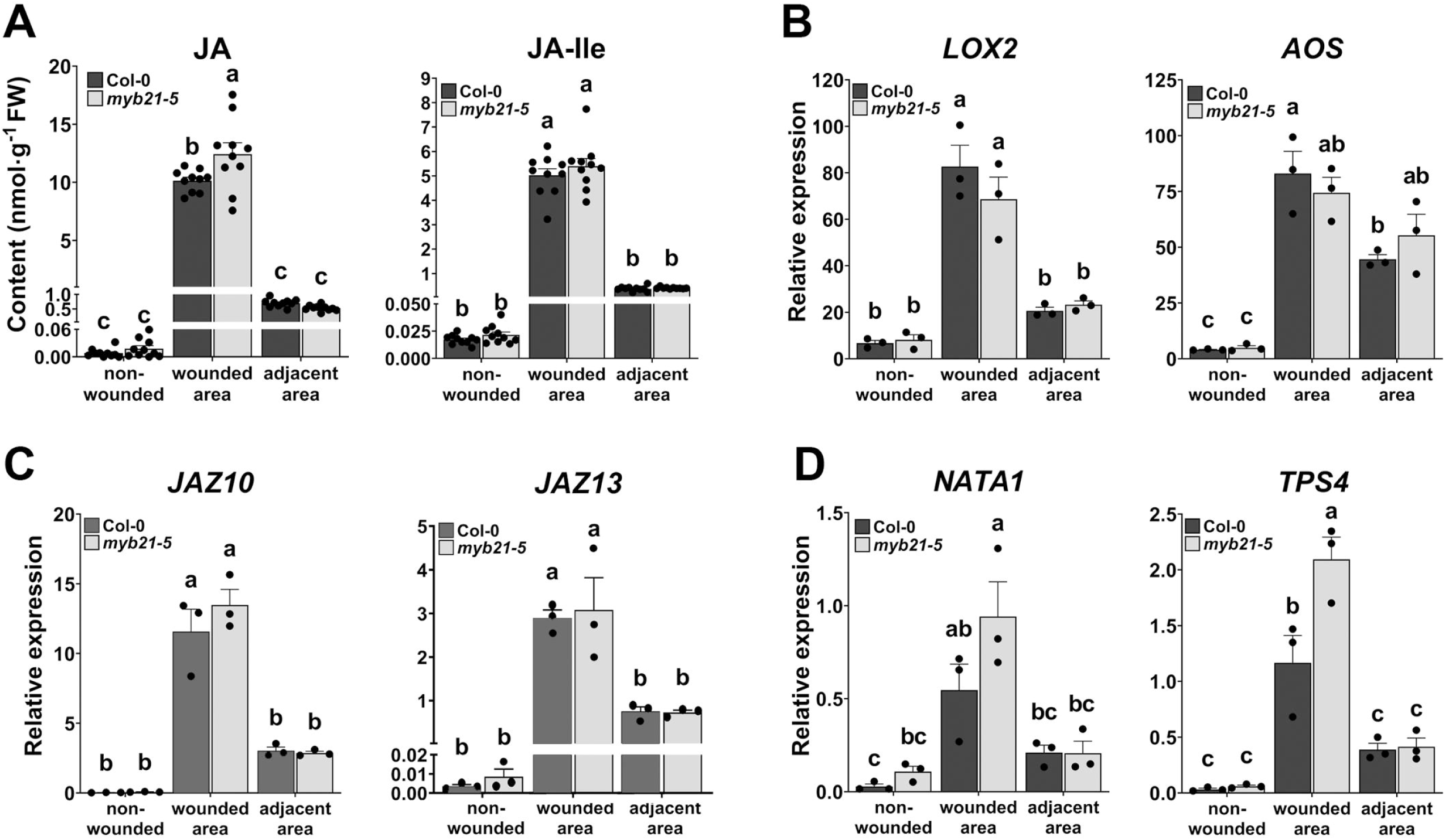
MYB21 does not regulate JA signaling and biosynthesis in the local or systemic wound-response of the leaf. Four-week-old leaves from *myb21-5* mutant and wild type (Col-0) plants were injured with a haemostat at the upper-half (wounded area) while the lower half remained non-wounded (adjacent area). Leaf halves were separated at 1 hour after wounding, with non-wounded leaves serving as controls. (A) Levels of JA and JA-Ile determined in relation to the tissue fresh weight. (B-D) Transcript accumulation of JA biosynthesis genes *LOX2* and *AOS* (B), JA signaling components *JAZ10* and *JAZ13* (C), and defense genes *NATA1* and *TPS4* (D), determined in relation to *AtPP2A3*. Bars represent means of ten biological replicates in A and three biological replicates in B-D (single dots, ±SEM). Letters denote statistically significant differences among samples as determined by two-way ANOVA followed by Tukey HSD (p<0.05). **Associated supplemental data: Figs. S3 and S4**

To gain further insight into a potential role of MYB21 in the JA-mediated response of plants, root growth inhibition by MeJA was tested using 10-day-old seedlings from wild type and *myb21-5* (Fig. 4). While *myb21-5* seedlings exhibited significantly longer root length compared to Col-0 seedlings at control conditions, MeJA inhibited primary root growth of both genotypes resulting in similar root length (Fig. 4A, B). This might suggest a higher root growth inhibition in *myb21-5* caused by a higher sensitivity of this mutant to MeJA. To test this at molecular level, transcript accumulations of *JAZ10* and *JAZ13* in seedlings were determined after treatment with 50 µM MeJA. There were no differences between wild type and mutant seedlings detected after 1 h of treatment (Fig. 4C). Treatment for 7 days resulted, however, in less transcript accumulation in mutant seedlings compared to wild-type seedlings (Fig. 4D). Both the longer seedlings roots under control conditions as well as the lower transcript accumulation of JA-induced genes after long-term treatment suggest that MYB21 might play a role in enhancing JA-responses resulting in growth repression during development and in response to increased hormone levels.

**Fig. 4:**
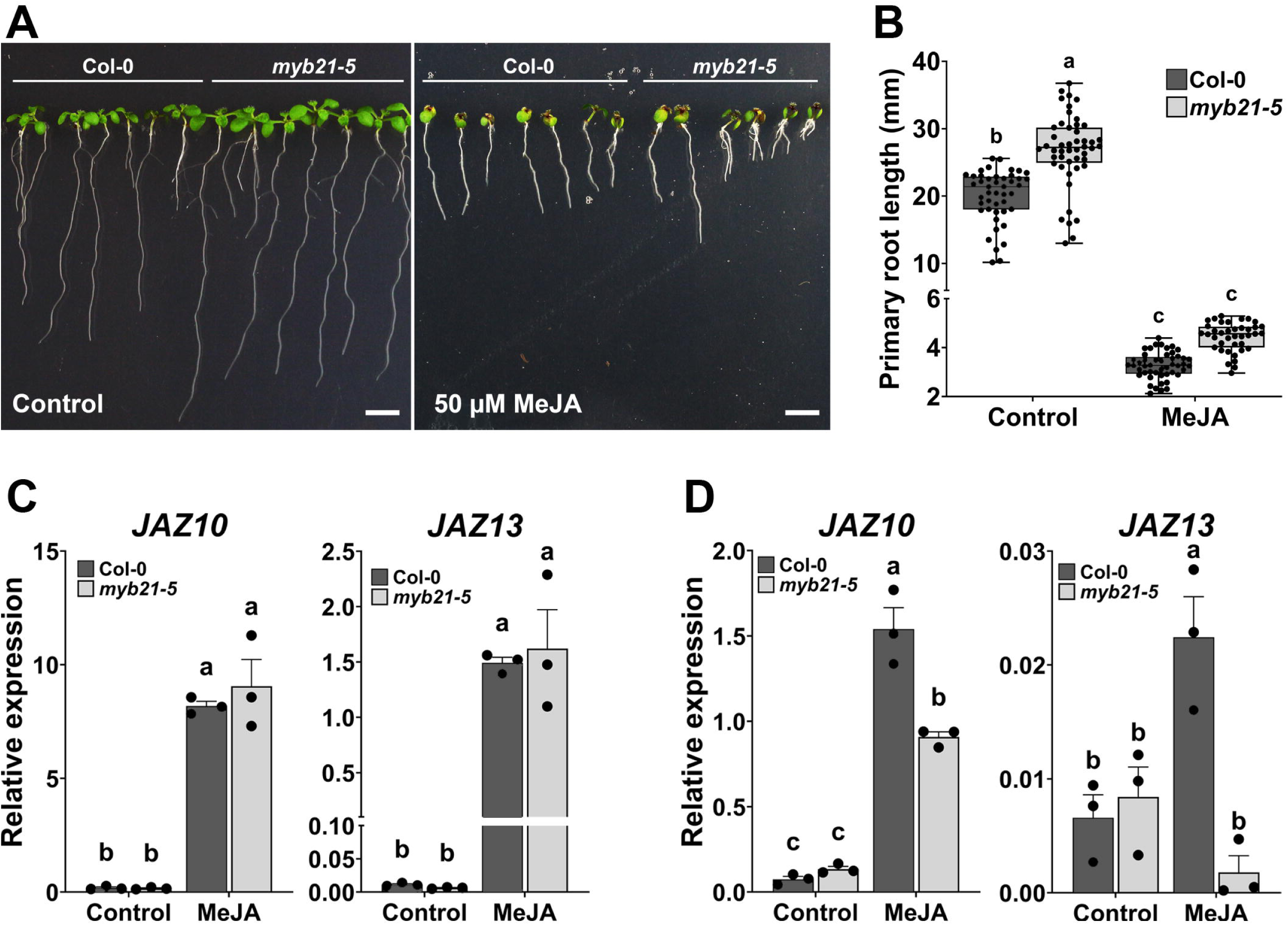
The *myb21-5* seedlings show wild type-like early response to MeJA but different late response. (A) *myb21-5* mutant and wild type (Col-0) seedlings grown in the absence (control) or presence of 50 µM MeJA. (B) Primary root length (control, n=48-45; MeJA, n=39-48) was measured 10 days after germination. Data are presented as box plots, with standard deviations and means depicted using solid lines, and individual dots representing the full data range. (C-D) Expression levels of the JA marker genes *JAZ10* and *JAZ13* in the seedlings determined at 1 hour (C) and at 7 days (D) after 50 µM MeJA treatment and calculated in relation to *AtPP2A3*. Bars are means of three biological replicates (single dots, ±SEM). Letters denote statistically significant differences among samples as determined by two-way ANOVA followed by Tukey HSD (p<0.05).

### The *myb21-5* mutant exhibits growth alterations indicating putative function of MYB21 in vegetative developmental processes

The *myb21-5* mutant plants grown under control conditions, either on soil or sterile media, produced larger plants compared to the wild-type Col-0. Fourteen days after germination, the leaves of the *myb21-5* mutant exhibited significantly greater leaf area, as well as longer petioles and leaf lengths compared to the wild type (Fig. 5A-C), despite both genotypes being at the same developmental stage of four developed leaves, corresponding to stage 1.04 as defined by (Boyes *et al*., 2001). The larger leaf area in the mutant correlated with a higher number of trichomes per leaf compared to the wild type (Fig. 5D). Consistently, the mutant primary root was significantly longer than the wild type in seven-day-old seedlings (Figs. 4A, B, 5E, F). Similar differences between *myb21-5* mutant and wild type were also observed in fully developed rosettes of five-week-old plants, where the rosette diameter in the mutant was significantly larger than the wild type (Fig. S5A-B). Given that *myb21-5* exhibited greater leaf area compared to wild-type, cross-sections from leaves eight and thirteen at fully developed rosette stage (four-week-old plants) were analyzed to assess cell size and number (Fig. 5G). The size of palisade parenchyma cells was approximately 25 % larger in the mutant leaf compared to wild type (Fig. 5G-H). This increase in cell size was observed only in leaf thirteen; palisade parenchyma cells in leaf eight did not differ significantly between mutant and wild type (Fig. 5H, Fig. S5C). Cell number, determined from the same sections, was similar between genotypes in both the adaxial epidermis and palisade parenchyma (Fig. 5I, Fig. S5C). Thus, the increased leaf area in the mutant is likely due to the enlarged size of individual cells rather than an increase in cell number. These vegetative growth alterations suggest a role of *MYB21* in repressing vegetative growth; a function known to be associated with JA (Noir et al. 2013; Staswick et al. 1992).

**Fig. 5:**
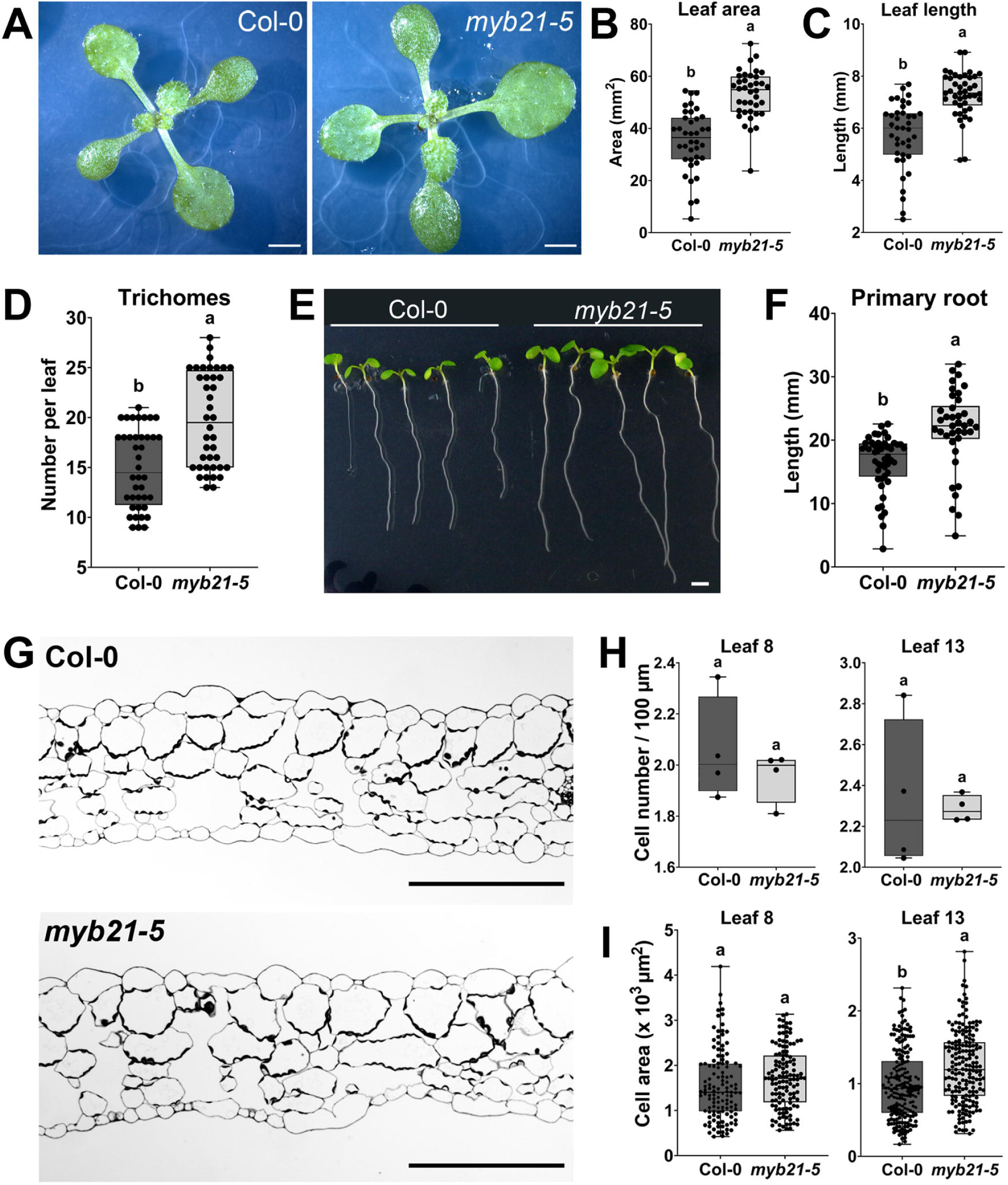
The *myb21-5* mutant exhibits vegetative growth defects compared to wild type (Col-0) under control conditions. (A) Representative pictures of plants of wild type and *myb21-5* at 14 days after germination, (B) leaf area, (C) length of the leaf, (D), trichome number per leaf. Note that *myb21-5* mutant shows increased values for all these parameters (n=38-40). (E-F) Primary root length at 7 days after germination (n=46-38). (G) Representative cross-sections of fully developed leaves of Col-0 and *myb21-5*. (H-I) Cell number and size of leaves No. 8 and No. 13 from wild type and *myb21-5*. Data are presented as box plots, with standard deviations and means depicted using solid lines, and individual dots representing the full data range. Letters denote statistically significant differences among samples as determined by Student’s *t*-test (p<0.05). Scale bars: 2 mm in A, 100 µm in G. **Associated supplemental data: Fig. S5**

In addition to vegetative growth, JAs are known to inhibit seed germination (Kumar *et al*., 2025). To evaluate whether MYB21 is involved in this process, rates of seed germination and development were determined in *myb21-5* and wild type (Fig. 6). Seed germination and seedling emergence occur within 72 hours after seed imbibition and comprise six stages, spanning from the stage of unbroken seed to fully emerged seedling with open cotyledons (Fig. 6A). Assessment of the seed germination rate was performed by counting the occurrence of each developmental stage (1-6) at different time points after imbibition.

**Fig. 6:**
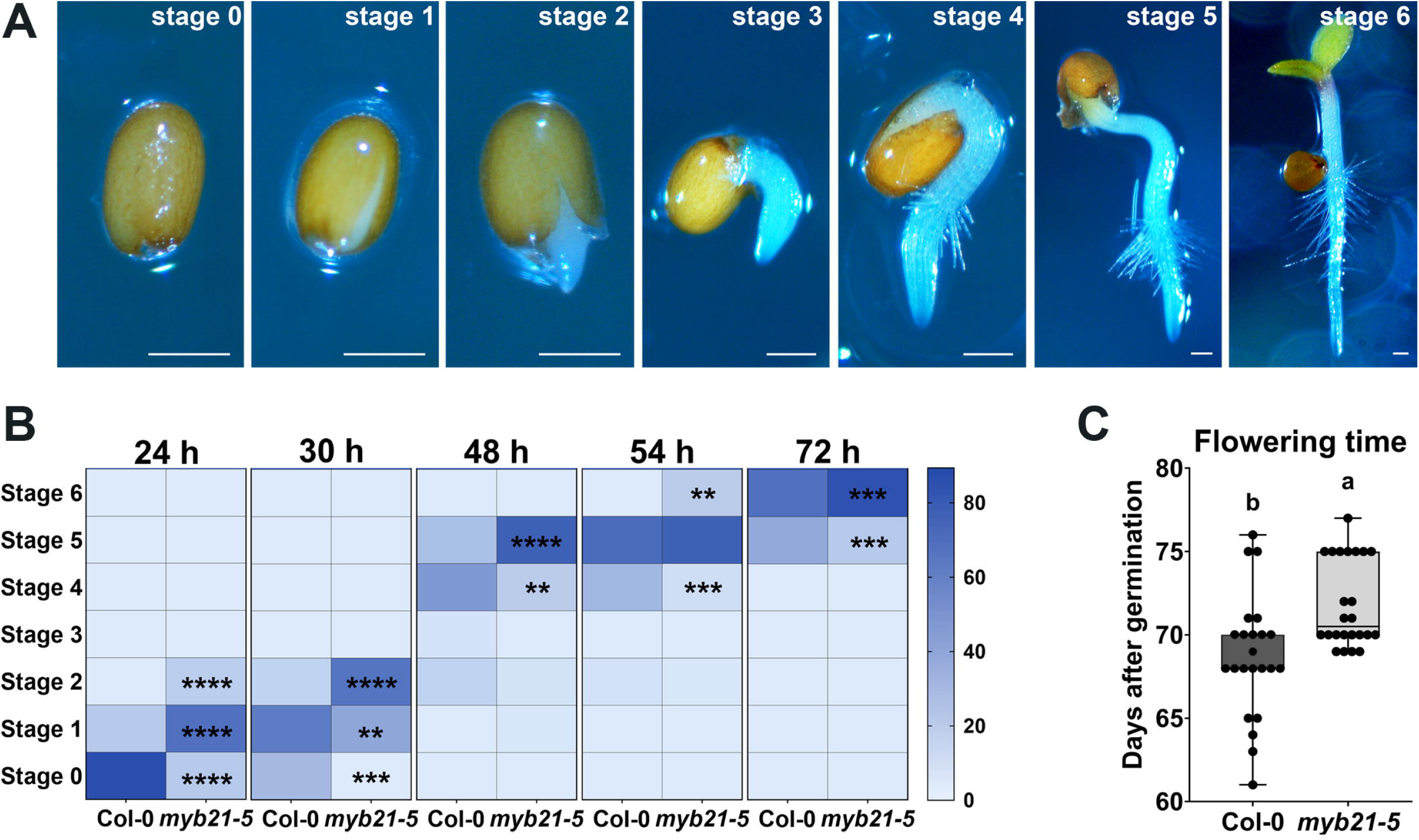
The *myb21-5* mutant germinates faster but flowers later than wild type. (A) Representative pictures of seed development stages with stage 0 = unbroken seed, stage 1 = testa rupture, stage 2 = radicle protrusion, stage 3 = primary root emergence, stage 4 = appearance of root hairs, stage 5 = whole root emergence and stage 6 = cotyledons emergence. (B) Quantification of each developmental stage (1-6) in the *myb21-5* mutant and wild type (Col-0) at 24, 30, 48, 54, and 72 hours after imbibition. (C) Bolting time of the *myb21-5* mutant and wild type (Col-0), determined as the number of days from germination to the elongation of the floral stem at 1 mm height (n=23-24). Data in (B) represents means of the percentage of seeds at each stage, calculated in relation to the total number of seeds (38-40 seeds per biological replicate; n=3). Data in (C) are presented as box plots, with standard deviations and means depicted using solid lines, and individual dots representing the full data range. Statistically significant differences among genotypes in (B) were determined by one-way ANOVA followed by Tukey HSD (p<0.05) and are indicated by asterisks (*p<0.05, **p<0.01, ***p<0.001, ****p<0.0001), and by Student’s t-test (p<0.05) in (C) indicated by letters.

Twenty-four hours after imbibition, 83% of the wild-type seeds were still at stage 0 of unbroken seed, while 63% of the *myb21-5* seeds were at stage 1 of broken testa, 19% at stage 2 and only 18% were still at stage 0 (Fig. 6B). *myb21-5* seeds showed accelerated germination rates most significantly between 24 and 48 hours after sowing leading to 78% fully emerged seedlings in the mutant and 55% in wild-type at 72 hours (Fig. 6B). These results indicate a role of MYB21 in the inhibition of seed germination. Since JA is a regulator of flowering time (Li *et al*., 2024), the flowering time (bolting time) of the *myb21-5* mutant and the wild type was also determined (Fig. 6C). Plants of *myb21-5* entered flowering stage with three days delay compared to wild-type, implying an involvement of MYB21 in Arabidopsis flowering time.

### Transcriptome analysis of the *myb21-5* mutant seedlings shows alterations in photosynthesis, lignin biosynthesis and defense responses

To elucidate the specific function(s) that MYB21 could exert in vegetative tissues, an mRNAseq analysis was performed on fourteen-day-old from wild-type and the *myb21-5* seedlings. The comparison between both transcriptomes resulted in 457 differentially regulated genes (DEGs) in *myb21-5* with a fold change (FC) ≥ 2, p < 0.05, and fragments per kilobase of transcript per million mapped reads (FPKM) ≥ 0.5 (Fig. 7A, Dataset S1). These DEGs showed KEGG enrichment in photosynthesis and phenylpropanoid-derived secondary metabolites including flavonoids and stilbenoid diarylheptanoid and gingerol (Fig. 7B). Similarly, gene ontology (GO) enrichment analysis revealed alteration of four main pathways in the *myb21-5* mutant corresponding to photosynthesis antenna proteins, lignin biosynthesis, defense response and response to cytokinin (Fig. 7C-F). For photosynthesis, four main genes encoding antenna proteins belonging to the photosystem II light-harvesting complex (LHCII), *LHCB2.3*, *LHCB2.2*, *LHCB4.2* and *LHB1B1*, were downregulated in the *myb21-5* mutant (Fig. 7C). These genes encode chlorophyll a/b binding proteins that form the antenna system of the photosynthetic apparatus (Guardini *et al*., 2025). The lignin biosynthesis branch of the phenylpropanoid pathway was also downregulated in the *myb21-5* mutant, with several genes involved in the formation of monolignols which are the aromatic alcohol precursors serving as building blocks of lignin (Fig. 7D), and comprised 4-coumarate:CoA ligases (*4CL1*, *4CL3*), *O*-methyltransferases (*OMT1*, *CCoAOMT1*), and four members of the cinnamyl alcohol dehydrogenase gene family (*CAD5*, *CAD7*, *CAD8* and *CAD9*). Additionally, key defense-related genes from the plant defensin family (PDFs), such as *PDF1.2*, *PDF1.2b*, *PDF1.2c*, and *PDF1.3*, showed lower transcript levels in the mutant compared to wild type (Fig. 7E). Alteration in cytokinin hormone signaling was also detected in *myb21-5* transcriptome and involved an upregulation of several members of the Type-A Arabidopsis Response Regulators (ARR) *ARR5*, *ARR6*, *ARR7*, *ARR9*, *ARR15*, which act as negative feedback regulators of cytokinin signaling (Fig. 7F). The reduced expression levels of plant defensins in the transcriptome of the *myb21-5* mutant, and specifically *PDF1.2* which is a marker of the jasmonate-dependent defense responses, suggests a role of MYB21 in JA-mediated defense.

**Fig. 7:**
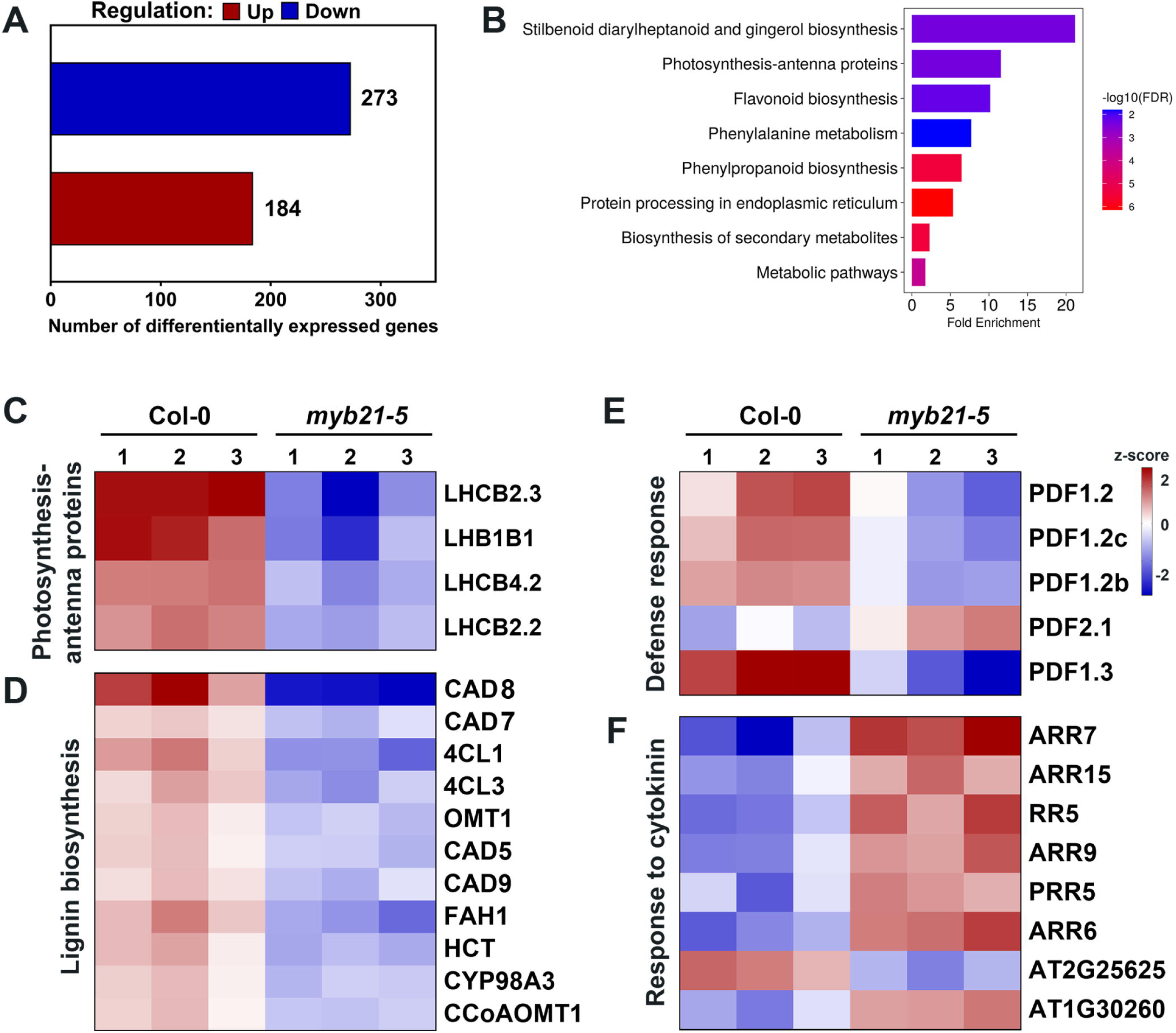
Transcriptome analysis reveals alterations in the biosynthesis of secondary metabolites pathway, photosynthesis, and defense genes in the *myb21-5* seedlings compared to wild type. (A) Differentially expressed genes (DEGs) in *myb21-5* seedlings compared to wild type (n=3), with a fold change (FC) ≥ 2, p < 0.05, and fragments per kilobase of transcript per million mapped reads (FPKM) ≥ 0.5. (B) Gene enrichment analysis of the DEGs using the KEGG database (false discovery rate (FDR) < 0.05). (C-F) Z-score-based heatmaps showing representative DEGs from enriched pathways according to Gene Ontology (GO) “Biological Processes” (BP) (FDR < 0.05). (C) Photosynthesis-antenna proteins. (D) Lignin biosynthesis. (E) Defense response. (F) Response to cytokinin. **Associated supplemental data: Dataset S1**

### *myb21* mutant plants show enhanced susceptibility to herbivore and pathogen attack

To investigate the possible contribution of MYB21 to the JA-mediated defense responses, we challenged the *myb21-5* mutant with insect feeding and pathogen assays and evaluated its performance in comparison to the *MYB21_pro_:MYB21* complementation line (Schubert *et al*., 2019) and the wild-type (Fig. 8). No major differences in rosette size were observed among genotypes grown for bioassay conditions of 8h light / 16h dark photoperiod (Fig. S6). Larvae of *Spodoptera littoralis* were allowed to feed on mature plants and consumed predominantly older leaves (Fig. S6). Remarkably, *S. littoralis* larvae feeding on *myb21-5* leaves gained significantly more weight than those feeding on wild-type plants or *myb21-5* plants complemented with *MYB21_pro_:MYB21* showing a mild, but significant effect of MYB21 in defense against insect herbivores (Fig. 8A). The *myb21* susceptibility phenotype was consistent also in necrotrophic pathogen assays with *Botrytis cinerea*, which caused larger lesions on *myb21-5* compared to wild-type leaves (Fig. 8B). Complementation of *myb21-5* with *MYB21_pro_:MYB21* resulted in a partial rescue of the lesion sizes only (Fig. S7). Taken together, these data show a functional link between MYB21 and JA-mediated defense in leaves.

**Fig. 8:**
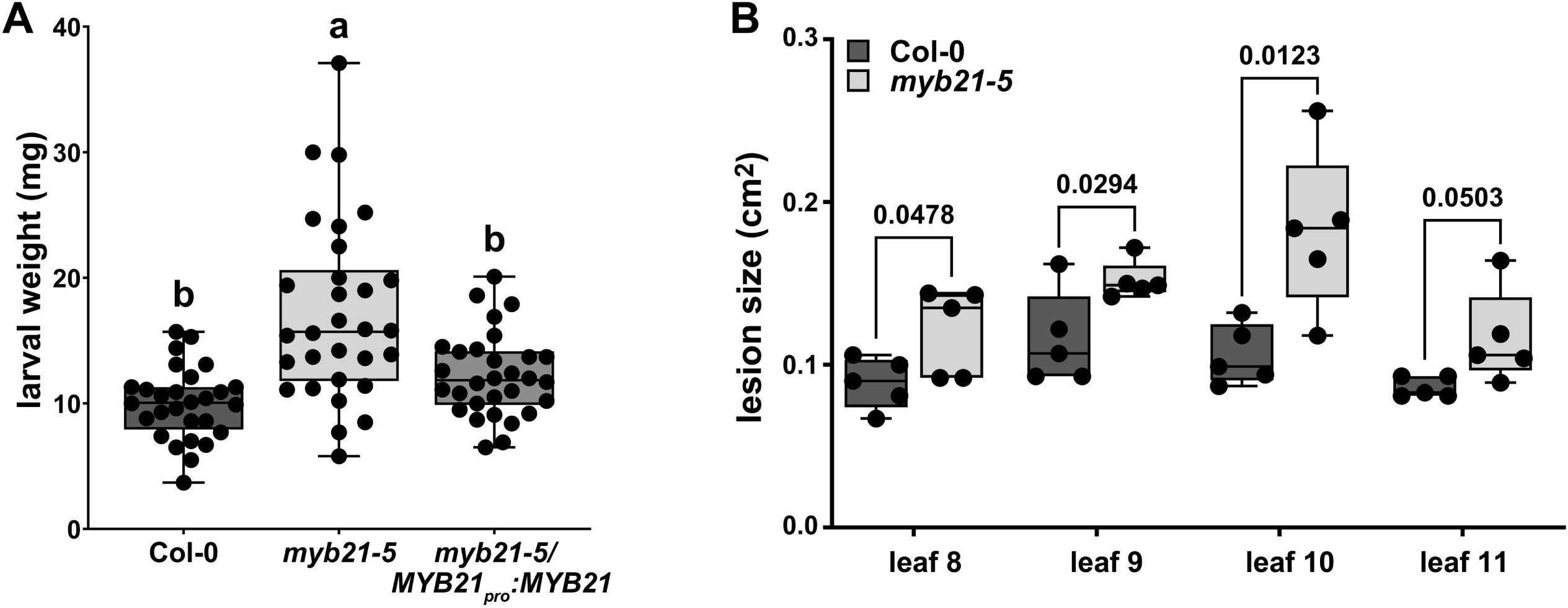
*myb21-5* plants show enhanced susceptibility to herbivores and pathogens. (A) Weight of *Spodoptera littoralis* larvae after feeding on five-week-old plants of the indicated genotypes for 9 days. Box plot summary is shown with medians represented by a solid line inside the box and circles showing individual data points. Letters denote statistically significant differences as determined by one-way-ANOVA followed by Tukey HSD (p<0.01). (B) Lesion size on wild-type (Col-0) leaves and leaves of *myb21-5* after infection with *Botrytis cinerea*. Leaves No 8-11 were evaluated separately and showed significant enlarged lesions for leaf No 8, 9 and 10 according to Student’s *t*-test. **Associated supplemental data: Figs. S6 and S7**

## Discussion

The R2R3-MYB transcription factor MYB21 is primarily known as a flower-specific transcription factor, and its function in stamen and flower development, as well as plant fertility has been characterized in Arabidopsis and tomato (Reeves *et al*., 2012; Schubert *et al*., 2019; Song *et al*., 2011). The detection of wound-induced *MYB21* transcript accumulation in Arabidopsis vegetative tissues led us to hypothesize that MYB21 may also exert functions in these tissues, despite its very low expression levels in non-floral organs.

During the transcriptional response to wounding, *MYB21* expression was induced in seedlings and mature leaves with a high fold-change but overall low transcripts levels. This wound-induced expression was JA-dependent, as it occurred in Col-0 but was abolished in the JA-deficient and JA-insensitive mutants *aos* and *coi1-16,* respectively. *MYB21* transcript accumulation was also responsive to exogenous JA treatment in leaves in a COI1-dependent manner, further indicating its JA-mediated regulation. In the absence of external stimuli, the almost undetectable levels of *MYB21* transcripts in the leaf and seedlings of Col-0 may result from repression of its expression in these tissues. Repression of *MYB21* expression in seedlings by the light-signaling component CONSTITUTIVE PHOTOMORPHOGENIC 1 (COP1) has previously been reported, as *MYB21* transcripts were detected in *cop1* mutant seedlings but not in wild-type plants (Shin *et al*., 2002). The stimulus responsible for the de-repression of *MYB21* in seedlings was, however, not identified. Here, we show that wounding and JA act as stimuli leading to *MYB21* transcript accumulation in leaves and seedlings and that increased *MYB21* expression clearly coincides with JA accumulation (Mekkaoui *et al*., 2025), JAZ degradation (Chung *et al*., 2008), and the induction of early JA-induced genes, such as *JAZ10* (Fig. 1). Together, these findings support a role for MYB21 in JA-mediated signaling in vegetative tissues, and specifically in the JA-mediated wound response. This is consistent with the previously characterized JA-dependent regulation of *MYB21* expression in flowers (Cheng *et al*.; Mandaokar *et al*., 2006; Reeves *et al*., 2012; Schubert *et al*., 2019).

Analysis of the *MYB21_pro_:GUSPlus* transcriptional reporter lines revealed a spatially restricted *MYB21* promoter activity to trichomes and hydathode areas in the absence of external stimuli. Upon JA treatment or wounding, induction of *MYB21* promoter activity remained highly cell-specific and was observed in the trichomes and their skirt cells, vascular tissues within the wounded area, and along the leaf margin.

Negative feedback regulation of JA biosynthesis by MYB21 has been reported in Arabidopsis flowers (Reeves *et al*., 2012). We were, however, unable to clearly confirm the presence of such a feedback-regulation role of MYB21 under either control conditions or following wounding in vegetative tissues. Although a slight increase in JA accumulation was observed at the wound site in *myb21-5* mutant leaves compared to wild-type plants, no statistically significant differences were detected in JA-Ile levels or the expression of JA marker genes. Instead, only minor upward trends in the expression of these marker genes were observed. These results suggest that MYB21-mediated feedback regulation of JA biosynthesis may be absent in vegetative tissues or that our experimental approach lacked sufficient sensitivity due to the spatially restricted expression of *MYB21* within the leaf, where high basal of JA biosynthetic enzymes throughout the tissue (Hause *et al*., 2003; Stenzel *et al*., 2003) may mask localized regulatory effects.

A possible role of MYB21 in JA-mediated root growth inhibition, leaf anthocyanin accumulation and plant defense against insect attack, was previously ruled out, as ectopic expression of *MYB21* in the JA-insensitive *coi1-1* mutant failed to restore JA-regulated root growth inhibition, anthocyanin accumulation, or plant survival under *Bradysia impatiens* attack (Song *et al*., 2011). However, our results revealed that MYB21 influences several growth and developmental processes that are known to be regulated by JA. The *myb21-5* mutant displayed alterations in several traits including leaf area, root length, trichome development and seed germination. Notably, the mutant exhibited significantly increased leaf area and root length, as well as accelerated early stages of seed germination, processes that are normally repressed by JA (Pan *et al*., 2020; Wasternack and Hause, 2013). In line with these observations, ectopic expression of *MYB21* in Arabidopsis has been reported to cause alterations in the vegetative phenotype, including severe dwarfism characterized by shorter stems and smaller leaves, reduced cell expansion in hypocotyls and seedling lethality (Shin *et al*., 2002).

Transcriptome analysis of *myb21-5* seedlings revealed reduced transcript levels of several lignin biosynthetic genes, suggesting a positive regulatory role of MYB21 in lignin biosynthesis in the leaf. Among these genes are *CAD5*, *CAD7*, *CAD8*, and *CAD9*, whose reported spatial expression patterns coincide with those of *MYB21* in leaf veins, trichomes and hydathode areas (Kim *et al*., 2007). This observation aligns with the established function of MYB21 in flowers, where it has been shown to promote lignin deposition and positively regulates the expression of lignin biosynthetic genes, including *CCoAoMT* and *CAD* (Zhang *et al*., 2021). In addition to lignin-related changes, *myb21-5* seedlings exhibited transcriptional alterations in the photosynthesis-related transcriptional program, primary cytokinin response markers, and defense-responsive genes, particularly defensins. These pathways have been associated with jasmonate signaling (Attaran *et al*., 2014; Jang *et al*., 2017; Vijayan *et al*., 1998). Collectively, these transcriptional alterations suggest that MYB21 may influence multiple regulatory networks in leaves, ranging from cell wall biosynthesis to defense responses. Consistent with the altered defense-related transcriptional profile observed in *myb21-5*, the mutant displayed increased susceptibility to the herbivore *S. littoralis* and to the necrotrophic pathogen *B. cinerea*. Although the susceptibility was significant, it was moderate, indicating that MYB21 contributes partially to defense responses against these attackers and acts at very local levels such as trichomes. The comparatively stronger role of MYB21 in floral tissues, together with its more subtle effects in vegetative organs, namely leaves and seedlings, may reflect tissue-specific functional specialization.

Together, these findings reveal a diversified role of MYB21 in leaves and seedlings of Arabidopsis, ranging from growth-related processes to defense, and extending beyond its well-established function in floral development.

## Supplementary data

**Fig. S1**: *MYB24* transcript accumulation in vegetative tissues upon wounding and MeJA treatment.

**Fig. S2**: Histochemical localization of *MYB21_pro_:GUSPlus* in a stage 13 flower bud and in leaves of 15-day-old seedlings.

**Fig. S3**: MYB21 does not regulate JA signaling and biosynthesis in wounded seedlings.

**Fig. S4**: Transcripts of *MYB21* accumulate in the local wounded but not in the systemic non-wounded area of a leaf.

**Fig. S5**: Diameter of rosettes from adult Col-0 and the *myb21-5* mutant plants.

**Fig. S6**: Representative plant phenotypes before and after insect feeding.

**Fig. S7**: Lesion size in *myb21-5* and its MYB21-complemented line after infection with *B. cinerea*.

**Table S1**: Primers sequences used for RT-qPCR

**Table S2**: *MYB21* expression in wounded Col-0 (RNAseq data)

**Dataset S1**: Differentially expressed genes in *myb21-*5 in comparison wild type

## Acknowledgements

We thank Hagen Stellmach (IPB Halle) for help in jasmonate quantification and Jason W. Reed (University of North Carolina) for providing seeds of *myb21-5*. Simone Fraas (Martin-Luther-University Halle-Wittenberg) is acknowledged for performing sectioning of leaves. We thank Edelgard Wendeler (MPIPZ Cologne) for technical support.

## Author contributions

KM, BH: conceptualization; KM, SGZ, GH, and DG: methodology; KM, SGZ, SM, and GH: formal analysis; KM, LTLN, OPH, SGZ, SM, RB, and GH: investigation; IFA and BH: resources; KM, SM, SGZ, and BH: data curation; KM and BH: writing - original draft; SGZ, SM, IFA, and DG: writing - review & editing; KM, SM, and BH: visualization; DG and BH: supervision; BH: funding acquisition.

## Data availability

The transcriptomic raw data generated in this study have been deposited in Sequence Read Archive (SRA) of NCBI (PRJNA1427093).

